# Shell-bound archives: uncovering nematode encapsulations in the Galapagos’ largest radiation

**DOI:** 10.1101/2025.11.03.686299

**Authors:** Alexandre Fuster-Calvo, Ian Oiler, Andrew MacDonald, Christine Parent, François Massol, Gustavo Burin, John G. Phillips, Robbie Rae, Dominique Gravel

## Abstract

How macroevolution interacts with ecological networks remains a major question in eco-evolutionary science. We investigate this interplay in the *Naesiotus* snail radiation of the Galápagos Islands, which encapsulates parasitic nematodes within the shell—a recently discovered gastropod defense. Using a natural history collection, we examined dry shells from 47 species across 12 islands, quantified encapsulations, and sequenced nematode DNA to reconstruct a host-parasite network. Encapsulations were widespread and revealed high nematode diversity, including in snail hosts presumed extinct. Nematode diversity was shaped by habitat, while encapsulation load was better explained by host species identity, suggesting species-specific defenses. Neither trait showed phylogenetic signal, and shell brightness was unrelated to nematode interactions. Similarly, host diversification rate did not predict network position, suggesting that macroevolution may leave a weak or obscured imprint on this host-parasite network. This snail-nematode system in islands readily enables integration of ecological networks, phylogeny, functional traits, and biogeography.

## Introduction

Understanding how macroevolutionary and ecological processes interact to shape biodiversity has long been a central goal in ecology and evolution (McGill et al. 2019). While species interactions underpin many major evolutionary theories—e.g., Red Queen dynamics (Benton 2009), ecological opportunity (Yoder et al. 2010), character displacement (Taper and Case 1992), coevolution (Thompson 2005)—we still lack a general framework explaining how network structure influences trait evolution and diversification at biogeographic scales, and vice versa (Martínez 2005; Allhoff and Drossell 2013; Harmon et al. 2019).

Recent integration of phylogenetic and ecological network data offers a path forward. Phylogenetic signal in network structure (Krasnov et al. 2012; Martín González et al. 2015; Aizen et al. 2016) and simulation studies exploring diversification modes and trait evolution (Braga et al. 2018; Hembry et al. 2018; Vitória et al. 2018) point to reciprocal feedbacks between macroevolution and interaction networks. Yet empirical systems linking these dimensions remain rare, due to the difficulty of collecting interaction, trait, and phylogenetic data across entire clades and biogeographies (McGill et al. 2019). Phylogeny construction demands extensive sequencing (Tang et al. 2015; Crampton-Platt et al. 2016), while interaction networks require intensive fieldwork across the full range of a clade (Rodriguez-Flores et al. 2019)—an effort rarely put together in practice.

Oceanic island radiations provide an exceptional opportunity to overcome these limitations. These radiations often involve striking ecological and morphological divergence within lineages, unfold in spatially discrete and temporally bounded settings, and are among the best-studied cases of diversification (Gillespie 2007; Losos and Ricklefs 2009; Warren et al. 2015). Many such lineages already have available phylogenetic and trait data (e.g., HervÍas-Parejo et al. 2019; Fernández-Mazuecos et al. 2020; but see Cerca et al. 2023), and their limited geographic and ecological scope facilitates clade-wide sampling of ecological interactions (Bellvert et al. 2023). Nonetheless, fieldwork across remote archipelagos remains logistically demanding (Heleno et al. 2013).

Parasitic interactions offer a promising solution (Runghen et al. 2021). They are central to coevolutionary theory (Hoberg and Brooks 2008; Nuismer et al. 2008; Kamiya et al. 2014; Buckingham and Ashby 2022), and offer practical advantages for empirical integration of ecological and evolutionary data: they are ubiquitous and influential ecological players (Dobson et al. 2008; Lafferty et al. 2008; Wood and Johnson 2015), and their small size and host association make them well suited for large-scale sampling and identification via DNA sequencing, enabling simultaneous reconstruction of host-parasite networks and phylogenies (Clare et al. 2019).

A recent discovery by Rae (2017) revealed an extraordinary adaptation: land snails trap, lyse, and permanently encapsulate nematodes in their shell matrix. This is the first known exoskeletal immune system, conserved across the Gastropoda clade, and highly effective—trapping hundreds of nematodes within days, from micrometer-long eggs to centimeter-sized adults (Rae 2017, 2018; Dahirel et al. 2022). Crucially, Rae (2017) demonstrated in *Cornu aspersum* specimens that it is possible to extract and amplify DNA from the encapsulated nematodes. This opens the possibility that similar analyses could be performed on dry shell collections, allowing accurate reconstruction of host-parasite interactions across time and space with minimal sampling effort, and eliminating the need to harm individuals, which is particularly important for the conservation of highly threatened insular faunas (Dumbacher and Chaves, 2023).

Snail shells, shaped by selection for predator avoidance and microclimate adaptation (Goodfriend 1986, Chiba and Cowie 2016), now emerge as a functional and evolutionary trait mediating parasitic defense. Because many land snails undergo adaptive radiation on oceanic islands (Chiba and Cowie 2016), this mechanism offers a unique opportunity to integrate ecological network and phylogenetic data across entire radiations, and compare studies across archipelagos.

### The *Naesiotus* Radiation

The genus of land snails *Naesiotus* (Bulimulidae) protagonizes the most species-rich adaptive radiation in the Galápagos Islands. Though ∼61 species have been formally described (Phillips et al. 2020), genetic evidence suggests many cryptic or extinct taxa, bringing the total to 80–90 species (CP and JP, unpublished). Species in the genus have colonized all major islands and many islets, with evidence for single-island colonization followed by in situ diversification within islands for several islands (Parent and Crespi 2006; Phillips et al. 2020).

On Galápagos, *Naesiotus* species diversity spans a wide range of habitats and exhibits marked variation in shell size, shape, color, and brightness—traits shaped by both thermoregulatory and predatory pressures (Kraemer et al. 2019). Shell brightness, in particular, may play an additional role in parasite defense, as pigments such as melanin are known to influence immune responses across taxa (Dubey and Roulin 2014), including land snails (Caglio et al. 2018). More broadly, we hypothesize that macroevolutionary dynamics may influence the structure of host-parasite interactions (and vice versa). Rapidly diversifying lineages—especially those shifting in habitats—may escape parasites, leading to more peripheral positions in interaction networks.

The adaptive radiation of Galápagos *Naesiotus* land snails offers a powerful system to explore the interface of diversification, trait evolution, and parasitism. Using a dry shell collection, we (1) quantify nematode encapsulations across *Naesiotus* species and islands; (2) identify encapsulated nematodes via DNA sequencing to reconstruct host-parasite networks; (3) assess how interactions relate to host traits, habitat, island characteristics, and phylogenetic relatedness; and (4) test whether diversification dynamics within *Naesiotus* correlate with network position.

## Methods

A key advantage of nematode encapsulation is that parasites remain embedded permanently in the snail shell, even in fossil specimens (Rae 2017). This allows the retrieval of host-parasite interactions from dry museum collections. We examined *Naesiotus* specimens from the Galápagos National Park collection, curated at the University of Idaho. For all analyses, we used the most recent time-calibrated *Naesiotus* phylogeny based on RADSeq data, which includes 74 terminal taxa, resolves previously ambiguous clades, and identifies cryptic species (Phillips et al., in prep; Figure 1b).

**Figure 1.**
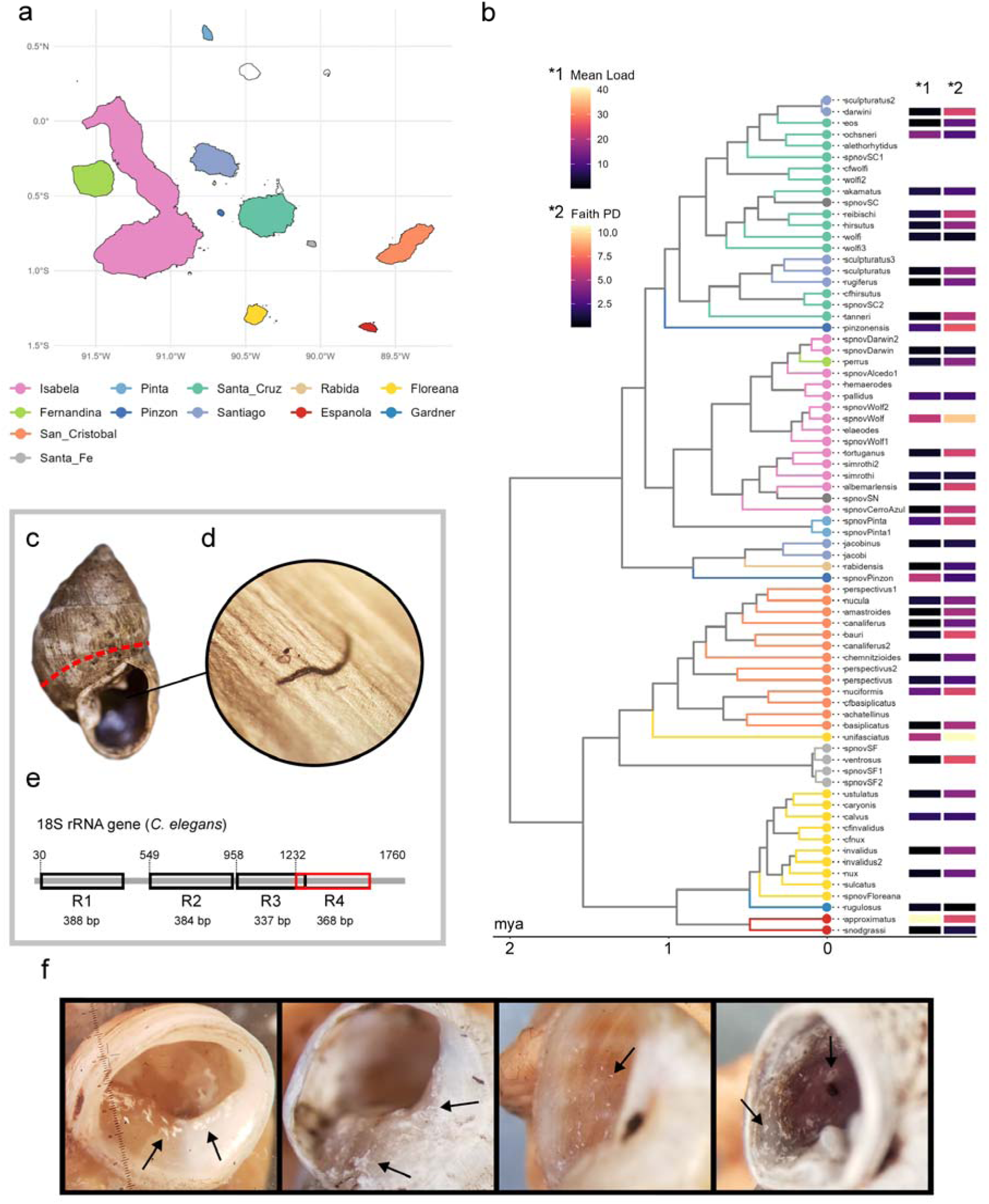
The *Naesiotus* radiation reveals a high diversity of encapsulated nematodes. Nematode encapsulations were observed in *Naesiotus* populations across the Galapagos archipelago. Champion Islet was sampled but grouped with Floreana due to its proximity and small size. (b) All 47 *Naesiotus* species examined showed encapsulations. Nematode DNA was successfully amplified in 46 species. Of these, 40 are represented in the phylogeny shown. The remaining 7 include six species considered extinct and one yet to be placed in the tree. Provisional names are used for new taxa for which no formal description has been made yet. Cryptic species are labeled with ‘cf.’ preceding the species name or with a numerical suffix, while clearly distinct undescribed species are denoted as “sp. nov. X”, where “X” refers to the island of origin. (c) Nematode load was quantified by counting visible encapsulations under an optical microscope on the external shell surface at the aperture, including the outer lip, palatal, parietal, and columellar walls. (d) Shell fragments containing visible encapsulations were sampled, (e) and nematode DNA was extracted and sequenced using primers targeting the fourth region of the 18S rRNA gene (schema adapted from Kenmotsu et al. 2020). (f) Examples of encapsulated nematodes in four different *Naesiotus* species.

From this phylogeny, we selected 47 species for encapsulation screening, determined by the availability of suitable specimens in the collection. For each species, we randomly selected at least 50 individuals and examined shells under a light microscope, recording the number of encapsulated nematodes visible in the adult shell aperture (outer lip, parietal, palatal, and columellar walls; Figure 1c–f). Loads exceeding 100 were recorded as “>100.” We avoided internal shell areas to prevent damage to specimens. For 32 species, sampling was increased to locate individuals with high encapsulation loads for DNA sequencing (Figure S1)

### DNA sampling and extraction

For each of the 47 species, we created positive and negative DNA pools. Positive pools contained shell fragments with visible encapsulations from 10 individuals (fewer when collection size was limited—average 9.26; range 1–10; Figure S2). Negative pools included 3 individuals without visible encapsulations, sampled from the entire aperture area. For two focal species—*N. unifasciatus* and *sp. nov. Wolf*—we performed individual-based sampling to assess completeness using rarefaction curves.

Fragments were crushed and decalcified with 0.5 M EDTA at 58°C overnight (Martin et al. 2021). DNA was extracted using the Qiagen DNeasy Blood & Tissue Kit. Sample volumes varied by shell size, and material was standardized by volume (∼0.2-0.5ml) per extraction tube.

### DNA sequencing and read processing

18S rRNA (V4 region) was amplified with primers NF1_MiseqF and 18Sr2b_ExtR_MiseqR (Kenmotsu et al. 2020) (see PCR conditions in Table S3) and sequenced using Illumina MiSeq (600-cycle v3 kit) at the CERMO-FC Genomics Platform, Université du Québec à Montréal. Read processing was conducted in QIIME2 2023.5 (Bolyen et al. 2019). DADA2 (Callahan et al. 2016) was used to generate ASVs with reads truncated at 220 bp. ASVs were clustered into OTUs at 97% identity with *vsearch* (Rognes et al. 2016). A phylogenetic tree of nematode OTUs was built using MAFFT alignment (Katoh & Standley 2013) and IQ-TREE (Minh et al. 2020) with ModelFinder (Kalyaanamoorthy et al. 2017), and used to compute Faith’s Phylogenetic Diversity (Faith 1992). In addition, taxonomy was assigned using a Naïve Bayes classifier trained on SILVA 138 (Quast et al. 2013) (see Supplement S1).

### Ecological and evolutionary predictors of nematode load and diversity

We used Bayesian hierarchical models to test whether nematode interactions reflect ecological predictors across *Naesiotus* species. Specifically, we modeled nematode encapsulation counts and phylogenetic diversity (Faith’s PD) as a function of snail shell brightness, snail microhabitat (arboreal vs. terrestrial), snail vegetation zone (arid vs. humid), and island characteristics (area and age). Trait data were compiled from Kraemer et al. (2019) and field observations, island area and ages from Geist et al. 2014 (Figure S6).

Because several species lacked complete trait data and island ages are uncertain, we added three other likelihood functions to our model. Shell brightness was modeled as a function of microhabitat, vegetation zone, island age, and area. Microhabitat (arboreal probability) and vegetation zone (arid probability) were modeled as binomial processes using species-level field count data where available. For species without field counts, these probabilities were estimated via partial pooling under hierarchical priors, sharing information across all species. Island age was included as a bounded latent variable drawn from uniform priors between minimum and maximum emergence estimates. This hierarchical approach allowed trait uncertainty and missing data to propagate through to the final predictions of nematode load and diversity.

Both models—encapsulation counts (Negative Binomial) and nematode diversity (Gaussian)—included species and island as group-level random effects. All models were implemented in Stan using the *cmdstanr* interface in R (Gabry and Cesnovar 2022). Details on model specification, parameters, and convergence diagnostics are provided in Supplement 1, section S1.2.

We tested for phylogenetic signal in both encapsulation load and nematode diversity by calculating Blomberg’s K and Pagel’s λ on the *Naesiotus* phylogeny using the *phylosig* function from the *phytools* R package (Revell 2024), with significance assessed via randomization-based tests.

### Relationship between diversification rate and network position

To test whether diversification dynamics correlate with host-parasite network structure, we estimated speciation rates for all terminal taxa in the *Naesiotus* tree using the ClaDS model (Maliet et al. 2019), which infers lineage-specific speciation rates under gradual heritable shifts at each branching event. The sampling fraction was 0.86 (73 species included out of an estimated total of 85).

We calculated three centrality metrics for each *Naesiotus* species in the regional host-parasite network: degree (interaction richness), betweenness (the extent to which a species lies on the shortest paths between other species, reflecting potential for mediating indirect effects), and eigen centrality (a measure of connectedness that gives more weight to links with well-connected partners) (Delmas et al. 2019). These were combined using principal coordinates analysis (PCA), and the first axis (explaining 90% of variance) was used as a composite centrality score (Figure S9).

We modeled centrality as a function of speciation rate, island age, island area (log-transformed), and local guild size (number of *Naesiotus* species per island). Predictors were standardized. To account for uncertainty in speciation rate estimates, we used 100 replicate models: in each replicate, we sampled one value from the posterior distribution of ClaDS speciation rates for each species, and refit the model. This generated a distribution of parameter estimates reflecting uncertainty in diversification rates. Each replicate was fit as a phylogenetic mixed model in MCMCglmm (Hadfield 2010), with a Gaussian error distribution and phylogenetic covariance matrix based on the *Naesiotus* tree. Species not present in the phylogeny or lacking nematode DNA data were excluded (n = 7). Details on model specification, parameters, and convergence diagnostics are provided in Appendix 5, section S5.3.

## Results

We analyzed 47 species from the *Naesiotus* radiation, examining 3,784 individuals collected from 247 localities across 10 main islands and two islets (Champion and Gardner). Champion was treated as part of Floreana island in the analyses due to its proximity and small size. On average, we sampled nematodes from 81 individuals (range: 17-178) and 5.7 localities (range: 1-33; Figure S1) per species. Six species were represented by fewer than 50 individuals due to miscounts or limited collection material. The most species-rich islands were Santa Cruz (10 species), San Cristóbal (9), and Isabela (7), while five islands and both islets each hosted a single *Naesiotus* species (Figure 1b, Figure 2).

**Figure 2.**
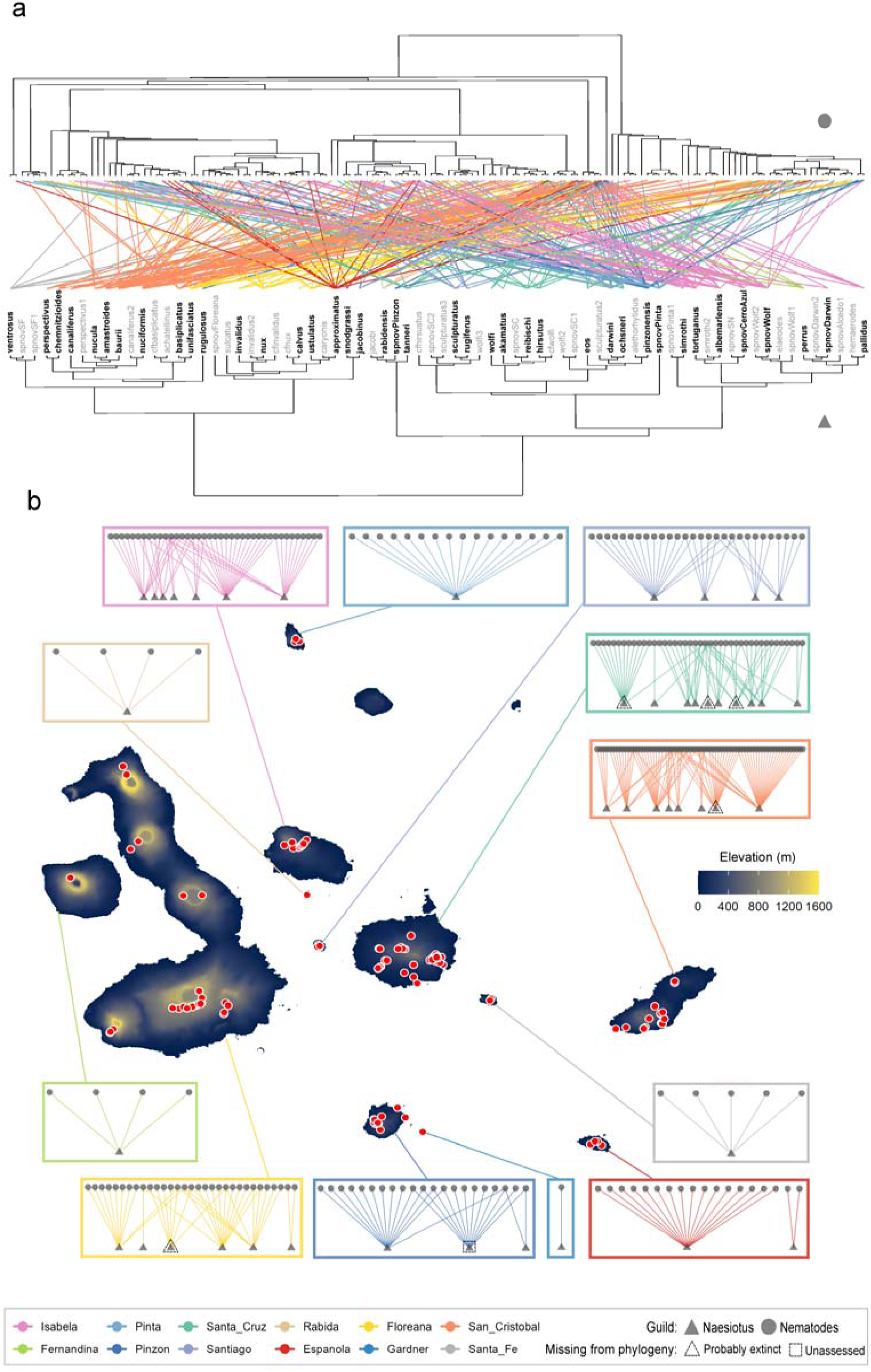
Naesiotus-nematode interaction network across the Galapagos archipelago. (a) A total of 147 nematode OTUs were identified from encapsulations in 47 *Naesiotus* species, revealing high parasite diversity across the radiation. The *Naesiotus* phylogeny is based on RADseq data, and the nematode phylogeny on the 18S rRNA gene. Species shown in black have been sampled. Seven species are not included in the tree: six are considered extinct, and one is pending assessment. Provisional names are used for new taxa for which no formal description has been made yet. Cryptic species are labeled with ‘cf.’ preceding the species name or with a numerical suffix, while clearly distinct undescribed species are denoted as “sp. nov. X”, where “X” refers to the island of origin. (b) Local interaction networks are shown for each island. Sampling sites (n = 247) are shown as red dots across the archipelago.

Encapsulation was found to be widespread, and nematode diversity high. Encapsulations were observed in all 47 analyzed species. The mean encapsulation load was 5.15 capsules (SD = 7.62; range: 0.13-40.9), and 42 species had at least one individual with more than 10 encapsulations (Figure 1b). On average, 70% of individuals per species had zero visible encapsulations in their shells (range: 13-99%).

We successfully amplified nematode DNA from 46 species, including all but sp. nov. Champion (Figure 2). This includes two species believed to be extinct, *cavagnaroi* and *lycodus*, with *cavagnaroi* hosting nematode OTUs unique to it (Figure S9). In addition, 25 species yielded nematode DNA from negative pools. In total, we identified 147 nematode OTUs representing 3 classes and 14 orders (Figure S5). Individual *Naesiotus* species hosted an average of 3.7 nematode orders (SD = 1.6; range: 1–7), with a mean Faith’s phylogenetic diversity (PD) of 4.25 (SD = 2.24; range: 0.06–10.7) (Figure 1b). Rarefaction curves did not reach an asymptote, indicating that the full diversity was not captured by our sampling (Figure S4). Neither encapsulation load nor nematode diversity showed significant phylogenetic signal across the radiation (Blomberg’s K = 1.1 and 0.5, P = 0.4 and 0.1; Pagel’s λ < 0.001, P = 1 for both; Figure S11).

Nematode encapsulation and diversity reveal distinct ecological signals. Shell brightness was strongly associated with microhabitat, with arboreal species being brighter, but it did not predict variation in nematode encapsulation or diversity (Figure 3a). Environmental predictors had a limited influence on encapsulation counts. Among fixed effects, only island area showed a weak positive effect. The estimated standard deviation of the species-level random effect was substantial (mean = 1.16, SD = 0.15, 95% CI: 0.89–1.48), indicating considerable interspecific variation in encapsulation load (Figure 3b).

**Figure 3.**
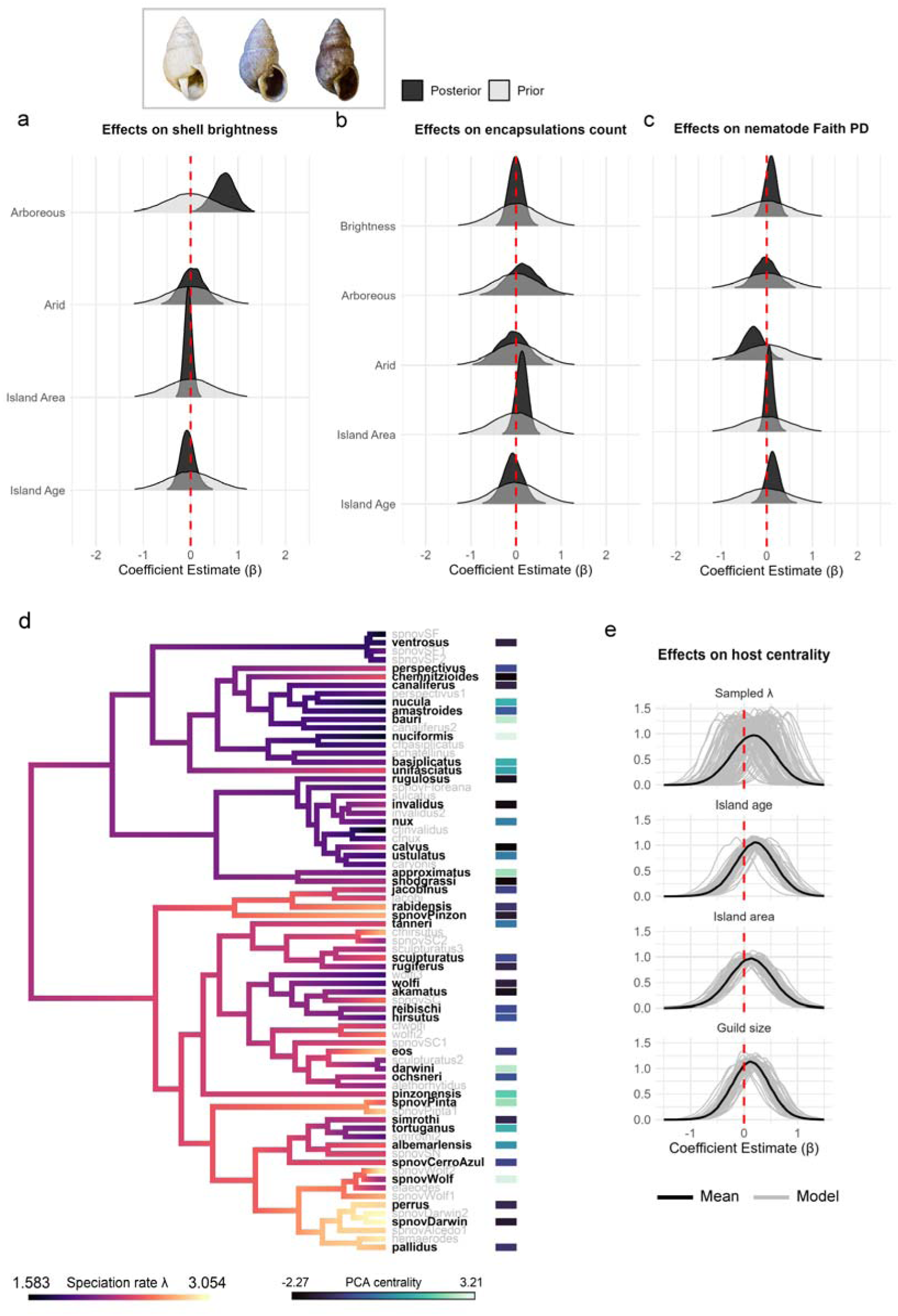
Posterior estimates linking shell traits to nematode load and diversity, and diversification rates to network position. Posterior distributions of models predicting (a) shell brightness, (b) encapsulation load, and (c) nematode diversity. (d) *Naesiotus* phylogenetic tree, with species included in analyses shown in bold. Speciation rates were estimated using ClaDS, and centrality scores correspond to the first principal component (PC1) from a PCA of degree, betweenness, and eigenvector centrality measures within the regional host–parasite network. (e) Posterior distributions summarizing the mean across 100 models, each sampling a speciation rate from the estimated posterior distribution of speciation rates.

In contrast, nematode Faith’s PD was significantly lower in species from arid vegetation zones. Island age and area showed weak positive associations with diversity. The species-level random effect for PD was smaller (mean = 0.20, SD = 0.13, 95% CI: 0.02–0.52), suggesting more moderate interspecific differences (Figure 3c).

We found no evidence that speciation rate predicts network position in the host-parasite interaction network. *Naesiotus* species’ centrality scores were not associated with speciation rate. Similarly, island area, island age, and guild size showed no clear effects on network centrality (Figure 3dc).

## Discussion

Our study presents a novel host-parasite system that integrates interaction networks, adaptive radiation, trait evolution, and island biogeography. By using nematode encapsulation in snail shells, we reveal widespread and diverse host-parasite associations across the largest adaptive radiation in the Galápagos, *Naesiotus*. Given the widespread occurrence of land snail radiations on both islands and continents, our findings open opportunities for comparative studies across archipelagos and between insular and continental systems. Such comparisons may reveal general mechanisms of coevolution-network interplay that help advance theory.

The consistent amplification of nematode DNA from shells without visible encapsulations indicates that our visual counts underestimate true infection rates and diversity. Some encapsulations may lie deeper within the shell matrix or form during earlier developmental stages, beyond the adult aperture we examined. Indeed, encapsulations extending toward the apex were occasionally observed. This raises the possibility of reconstructing temporal networks over the ontogeny of the hosts, or across seasonal and interannual variation (e.g., El Niño and La Niña), using preserved museum material.

We observed no phylogenetic signal in nematode encapsulation load or diversity, suggesting that parasitism is not phylogenetically conserved within *Naesiotus*. We also did not detect strong environmental effects on encapsulation load, suggesting that exposure alone does not explain variation in host–parasite interactions. If it did, we would expect higher loads in humid, vegetated habitats—conditions favorable to nematodes (Van Den Hoogen et al. 2019; Xiong et al. 2020). Instead, species identity emerged as the dominant predictor, highlighting the likely role of intrinsic traits such as immune defenses or life-history strategies. In contrast, nematode diversity suggested environmental and biogeographic filtering, with lower diversity in arid habitats and slightly higher on larger, older islands.

Notably, shell brightness, shaped by thermoregulation and visual crypsis (Kraemer et al. 2019), showed an apparent positive association with nematode diversity despite no effect on load. This may reflect increased exposure among brighter individuals in humid zones, where higher foraging activity (Kraemer et al. 2019) could lead to encounters with more diverse nematode lineages. In contrast, encapsulation load may remain unaffected if immune defense efficiency is unrelated to shell pigmentation, as was also found in Cepaea nemoralis (Williams & Rae 2016; Dahirel et al. 2024).

Mechanistically, we lack a clear understanding of how nematodes interact with the host. Although some may be commensals or passively encountered, others are likely true parasites (Barker 2004; Rae 2017). Chemical cues on the nematode cuticle may trigger targeted immune responses (Lee 2002), but the specificity and function of this defense remain open questions. Crucially, we cannot know whether higher encapsulation counts may reflect greater environmental exposure, differences in general immune capacity, or specific immune recognition (i.e., encapsulated nematodes might be those less able to evade host detection). Controlled experiments, though ideal, may be infeasible in threatened insular lineages like *Naesiotus*. Future studies could quantify nematode abundance in soil to test environmental exposure and examine how encapsulation load scales with snail age. Linear age-load relationships would indicate accumulation driven by exposure, while immunity differences might lead to disproportionate loads in older individuals.

The lack of association we found between speciation rates and network centrality of *Naesiotus* suggests that macroevolutionary dynamics may be decoupled from present-day interaction networks, or that other factors modulate the relationship. For example, centrality may depend more strongly on ecological traits such as population density or range size, which influence encounter rates and parasite transmission (Dallas et al. 2019). It is also possible that diversification and parasitism interact in more context-dependent ways: in some cases, parasites may be less able to track hosts that shift into novel microhabitats (Hoberg & Brooks 2008), while in others, strong parasitism pressure could inhibit diversification by imposing ecological constraints (Hasik et al. 2025). Uncertainty in our speciation rate estimates may have also limited our ability to detect clear relationships. While we inferred host diversification from a robust time-calibrated phylogeny, the nematode side remains poorly resolved. Expanding our preliminary 18S-based tree into a full phylogenomic reconstruction would be a critical next step to test diversification-centrality relationships within nematodes and the detection of co-speciation or host-shift dynamics (Fountain-Jones et al. 2018).

In terms of conservation, our work makes a major contribution to the poorly known nematode fauna of the Galápagos. Before this study, only 14 nematode species had been reported from two islands (Herrera et al. 2014), with a handful of earlier records (Buisán 1977; De Ley & Coomans 1990; Abebe & Coomans 1995). We now document 147 OTUs spanning 14 orders, including lineages also found in Galápagos tortoises (Fournié et al. 2015) and endemic cockroaches (Sinnott et al. 2015). Nematodes are, in fact, likely widespread across animal and plant hosts in Galápagos food webs (Lafferty et al. 2008). Understanding their life cycles and ecological interactions will be essential for conservation, especially as native hosts with reduced parasite loads may be particularly vulnerable to novel introductions (Parker et al. 2006). Network-based approaches can help detect these emergent threats and inform proactive management (Mougi 2022).

Finally, our study underscores the invaluable role of natural history collections in eco-evolutionary research (Meineke et al. 2019, Goulding et al. 2021). Because nematode encapsulations persist for decades or centuries, even in fossil shells (Rae 2017), they offer a unique opportunity to reconstruct historical host-parasite networks. Remarkably, we detected encapsulations in five *Naesiotus* species considered extinct, including nematode OTUs not found in any extant species. This suggests that these extinctions have led to undocumented losses in parasite diversity, potentially altering the structure and dynamics of parasitic networks or soil food webs (Lafferty 2012; Dallas & Cornelius 2015).

Our findings demonstrate that nematode encapsulations open the door to cost-effective network assessment of eco-evolutionary dynamics across space, time, and environmental change. More broadly, this work shows that sampling host-parasite interactions across adaptive radiations holds significant potential for advancing theory and contributing to the future development of eco-evolutionary science under a network perspective.

## Acknowledgements

We thank the Galápagos National Park Directorate for field and logistical support, and for facilitating the collection and export permits for samples used in this study. We also thank Jane Dostart and Kelly Martin for their assistance in managing the collection and supporting the DNA extraction protocols. We thank Geneviève Bourret for her contributions to the sequencing protocol and for leading the sequencing process. We also thank Andrew Kraemer for assisting with the shell brightness data and Guillaume Blanchet for helpful feedback on the statistical models. Lastly, we thank the Quebec Centre for Biodiversity Science (QCBS) and the Computational Biodiversity Science and Services (BIOS²) training program for supporting AFC’s research stays at the University of Idaho. Financial support was provided by the NSERC - CREATE Training program in computational biodiversity science and the NSERC Discovery Grant to DG.

## Notes

### Competing Interest Statement

The authors have declared no competing interest.

