## Supplement 1 for "Shell-bound archives: uncovering nematode encapsulations in the Galapagos’ largest radiation"

### S1.1 Sampling

**Table S1. PCR specifications.**

| Mastermix components | Final concentration | PCR program | |
|---|---|---|---|
| Molecular Grade H20 | n/a | Temperature °C | Time (minutes) |
| 5X Phusion HF Buffer | 1x | 98 | 0:30 |
| dNTPs mix 10 mM | 0.2 mM | 98 | 0:15 |
| DMSO | 3% | 58 | 0:30 |
| Primer NF1_F | 0.4 uM | 72 | 1:00 |
| Primer 18Sr2b | 0.4 uM | 72 | 10:00 |
| Phusion Hot Start II DNA pol. (2U/ul) | 0.02 U/ul | 8 | Hold |

*Add 1ul of DNA template diluted 1/10

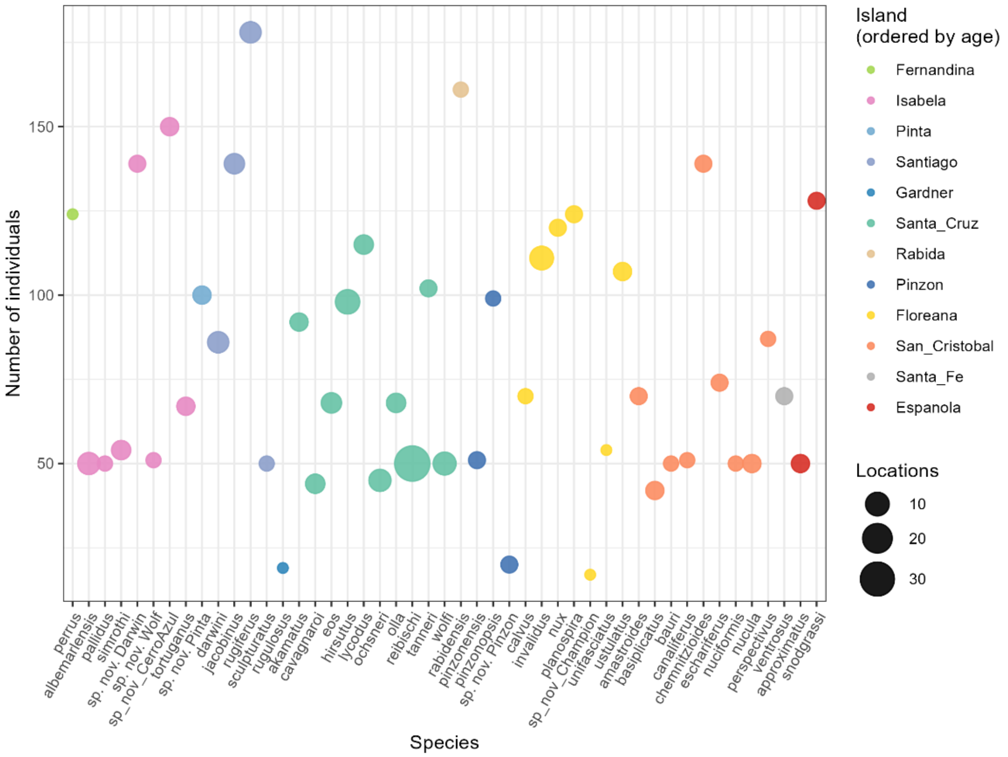


**Figure S1. Nematode encapsulation counts for the 47 studied *Naesiotus* species across Galápagos Islands.**

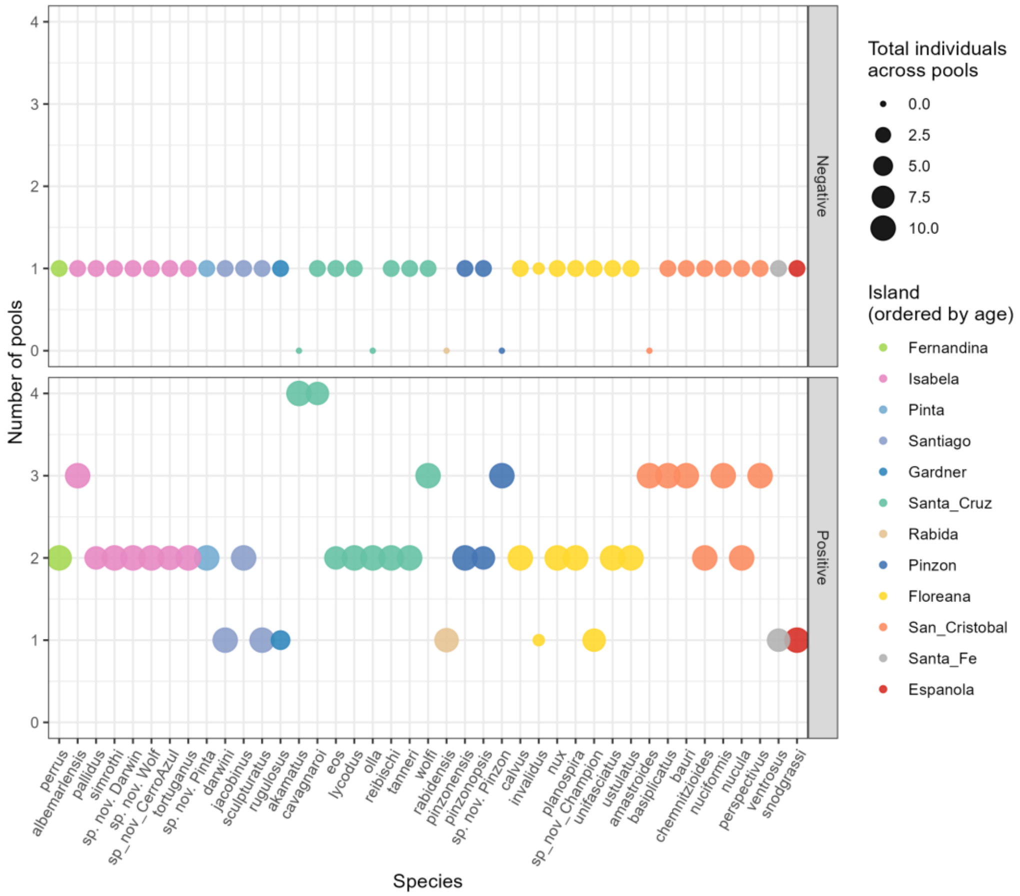


**Figure S2. Number of individuals in sequencing pools for the 47 studied *Naesiotus* species across Galápagos Islands.**

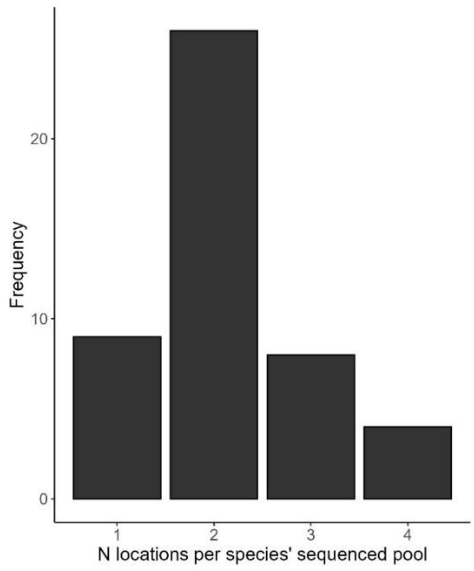


**Figure S3. Number of different locations of individuals per *Naesiotus* species' sequenced pools.**

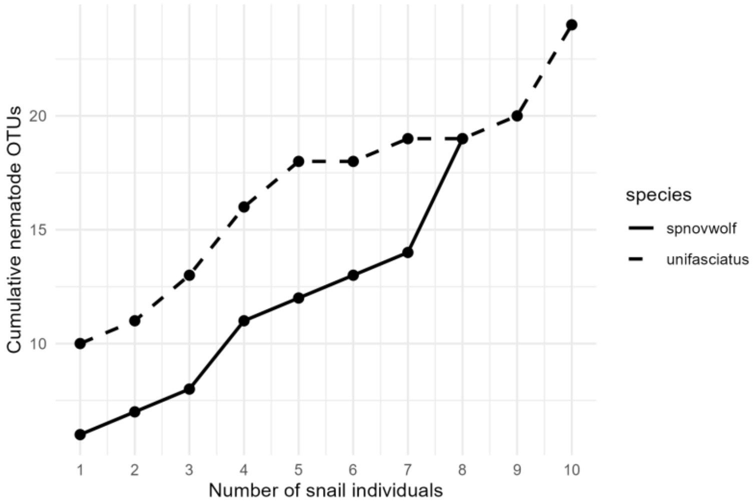


**Figure S4. Rarefaction curves assessing sampling completeness for nematode diversity in *Naesiotus* species.**

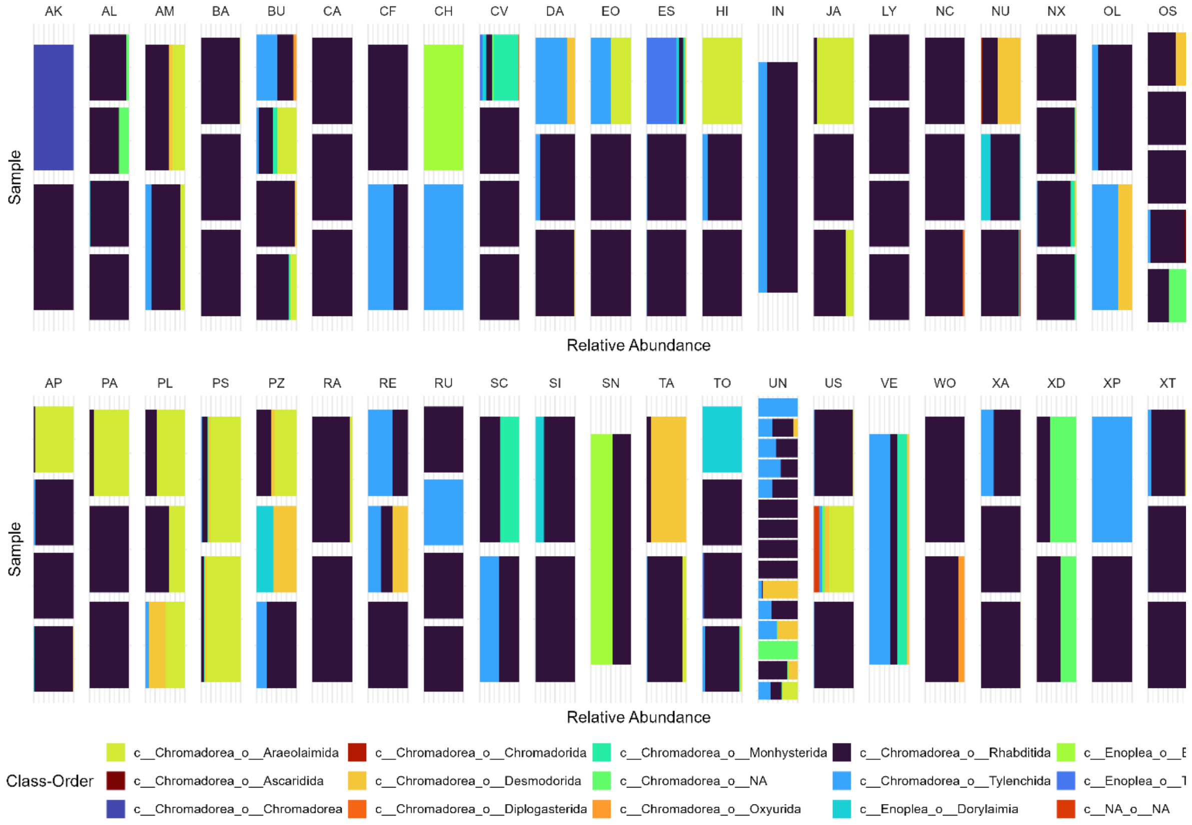


**Figure S5. Nemaode taxonomy assigned to OTUs in each sequenced *Naesiotus* species sample.**

Each sample is derived from pooled shell fragments containing encapsulations. On average, 9.26 snail individuals were pooled per species, and each sample contains shell fragments from a mean of 4.31 snails.

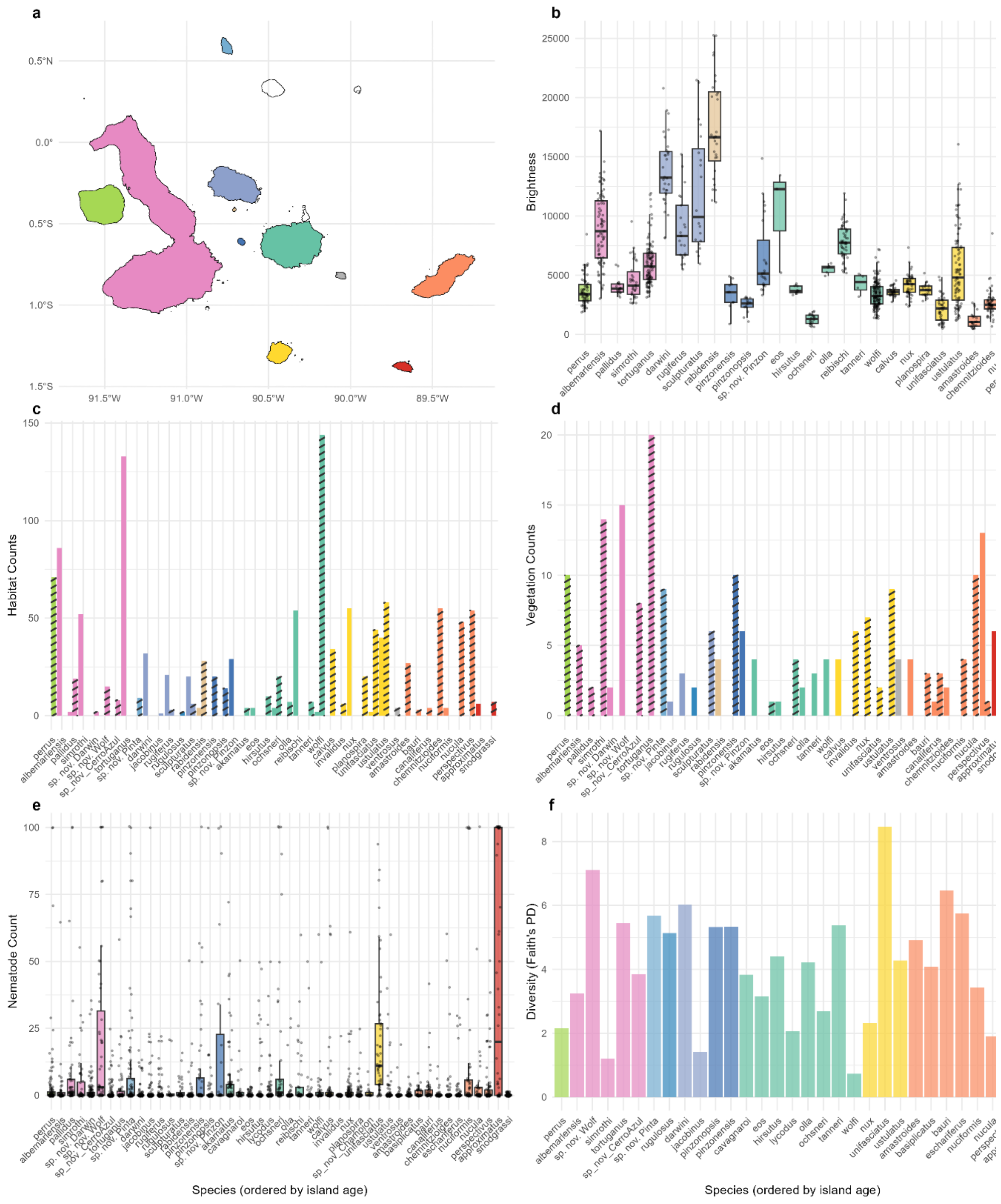


**Figure S6. Data on *Naesiotus* species' shell brightness, habitat, nematode load (number of encapsulations), and nematode diversity.**

### S1.2 Ecological predictors of nematode encapsulation and diversity

In the *Naesiotus* radiation, shell brightness appears to mediate a trade-off between abiotic selection for thermoregulation and biotic selection by predators. Kraemer et al. (2019) demonstrated that brightness increasingly matches background substrate as island age increases, suggesting that coevolution with predators like the endemic Galapagos mockingbird (*Mimus parvulus*) may drive biotic selection. They also found that background matches in brightness vary by microhabitat and vegetation zone.

We hypothesize that the adaptive variation in shell brightness shaped by thermoregulatory and predatory pressures may also influence host-parasite interactions with nematodes (Scheil et al. 2014). While the biochemical mechanisms underlying shell pigmentation remain poorly understood (Williams 2017), pigments like melanin are known to provide protection against both abiotic UV radiation and parasites across taxa (Dubey & Roulin 2014). Unlike previous work by Kraemer et al. (2019), which focused on background matching as a proxy for visual crypsis, our interest lay in the physiological and potentially immunological implications of pigmentation. We therefore used shell brightness itself (reflectance from 300 to 700 nm) as the focal trait.

To explore this, we used individual-level data on *Naesiotus* shell brightness (Kraemer et al. 2019), vegetation zone (humid or arid), and microhabitat (arboreal or terrestrial) (Figure S4). We then fit two Bayesian hierarchical models to examine how these traits, along with island characteristics (age and area), influence the number of encapsulated nematodes and their phylogenetic diversity (Faith's PD). Both models shared the same structure and incorporated species and island-level random effects.

The overall model for nematode load is structured as follows:

$$\log(Encapsulations_s) \sim \mu_{load} + \beta_{bright} log(Brightness_s) + \beta_{arbor} + \beta_{arbor,s} + \beta_{arid} + P_{arid,s} + \beta_{age} Age_i + \beta_{area} \log(Area_i) + u_{sp[s]} + u_{island[i]} \quad \text{Equation S1.1}$$

$$Encapsulations_s \sim NegBinomial(Encapsulations_{sp[n]}, \phi)$$

And for nematode diversity:

$$\log(\mu_s) \sim \beta_0 + \beta_{FPD} FPD_s + \beta_{bright} \log(Brightness_s) + \beta_{arbor} P_{arbor,s} + \beta_{arid} P_{arid,s} + \beta_{age} Age_i + \beta_{area} \log(Area_i) + u_{sp[s]} + u_{island[i]} \quad \text{Equation S1.2}$$

$$FaithPD_s \sim Gamma(\mu_s, \phi)$$

Where $Brightness_s$, $P_{arbor}$, and $P_{arid}$ are three species-level traits: shell brightness, and proportions of occurrences in arboreal or arid vegetation, respectively. $Age_i$ is the island age, $u_{sp}$ and $u_{island}$ are species and island random effects, and $\phi$ is a dispersion parameter.

We are missing species information for $Brightness_s$, $P_{arbor}$, and $P_{arid}$; in addition, island ages are only estimated within a broad range. To include all this uncertainty, we added four other likelihood functions to our model; this allows us to measure species traits and island ages as latent variables.

Brightness:

$$\mu_s \sim \beta_0 + \beta_{arbor} P_{arbor,s} + \beta_{arid} P_{arid,s} + \beta_{age} Age_i + \beta_{area} \log(Area_i) + u_{sp[s]} + u_{island[i]} \quad \text{Equation S1.3}$$

$$\log(Brightness_s) \sim Normal(\mu_s, \sigma^2)$$

Micro-habitat (terrestrial/arboreal):

$$Arboreal\ count_s \sim Binomial(N_{hab,s}, P_{arbor,s}) \qquad \text{Equation S1.4}$$

with $logit(P_{arbor,s}) \sim N(\mu_{arbor}, \sigma_{arbor})$

And for vegetation zone (humid/arid):

$$Arid\ zone\ count_s \sim Binomial(N_{veg,s}, P_{arid,s}) \qquad \text{Equation S1.5}$$

with $logit(P_{arid,s}) \sim N(\mu_{arid}, \sigma_{arid})$

And island age:

$$Age_i \sim uniform(min.emergence_i, max.emergence_i) \qquad \text{Equation S1.6}$$

$N_{hab,s}$ is the total number of habitat observations for species *s* (terrestrial or arboreal), and $N_{veg,s}$ the total number of observations of vegetation zone (humid or arid). For species missing microhabitat or vegetation zone data ($N_{hab,s}$ or $N_{veg,s}$ = 0), $P_{arbor,s}$ and $P_{arid,s}$ are treated as latent variables drawn from the hierarchical model.

This allowed uncertainty in trait estimates to propagate through to the final predictions. Models were specified in Stan and executed via the *cmdstanr* interface in R (Gabry et al. 2025). For the nematode load model, we used 10 parallel chains with 1000 post-warmup iterations each (10,000 posterior samples total). For the diversity model (Faith's PD), which required more complex sampling, we ran 6 parallel chains with 6000 post-warmup iterations each (36,000 posterior samples total), using an increased adapt_delta of 0.999 and max_treedepth of 18 to improve convergence.

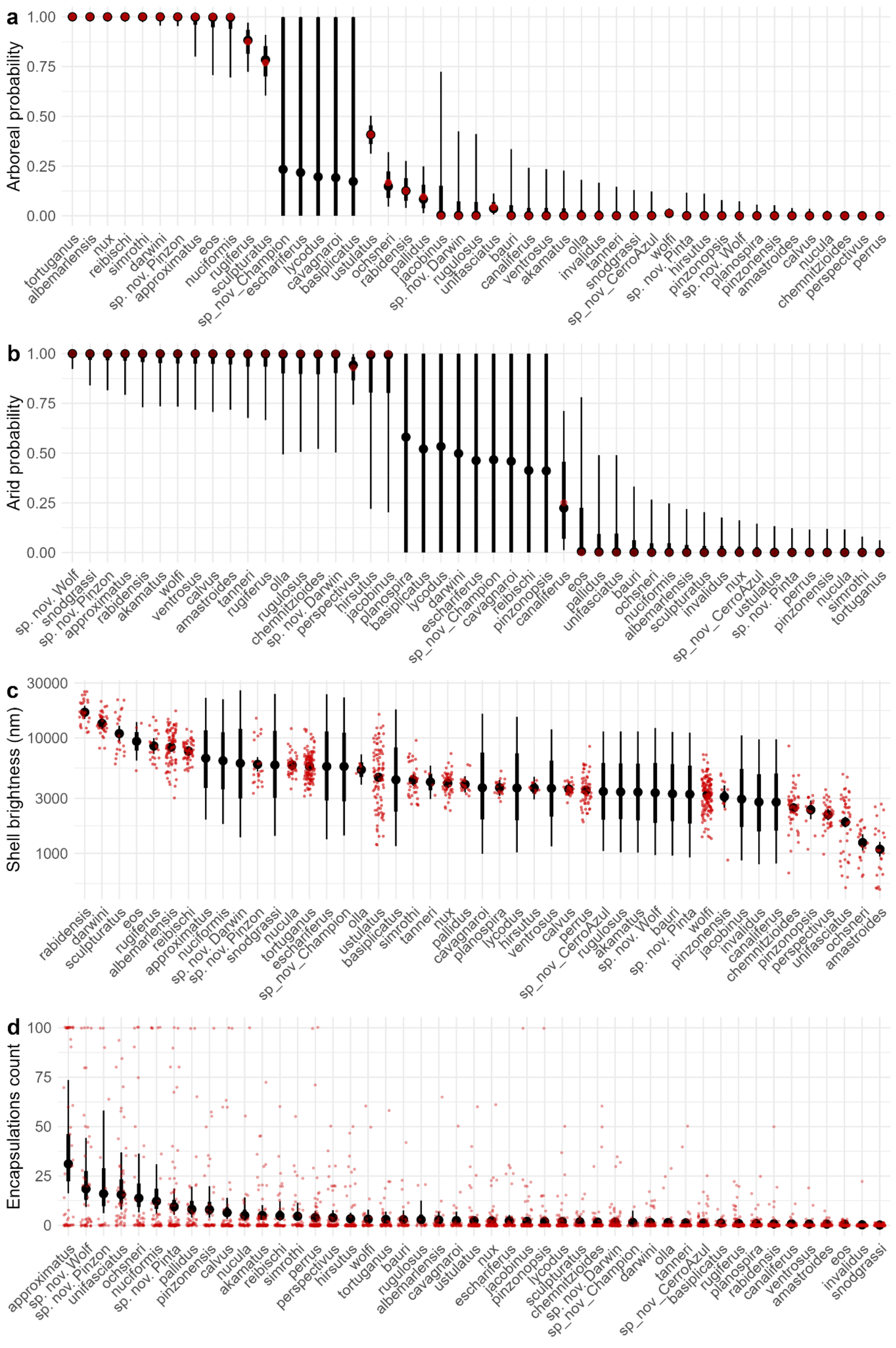


**Figure S7. Model predictions for encapsulation load (Model Equation S1.1).**

Red points show the raw data; black points show model predictions (posterior means with 95% credible intervals).

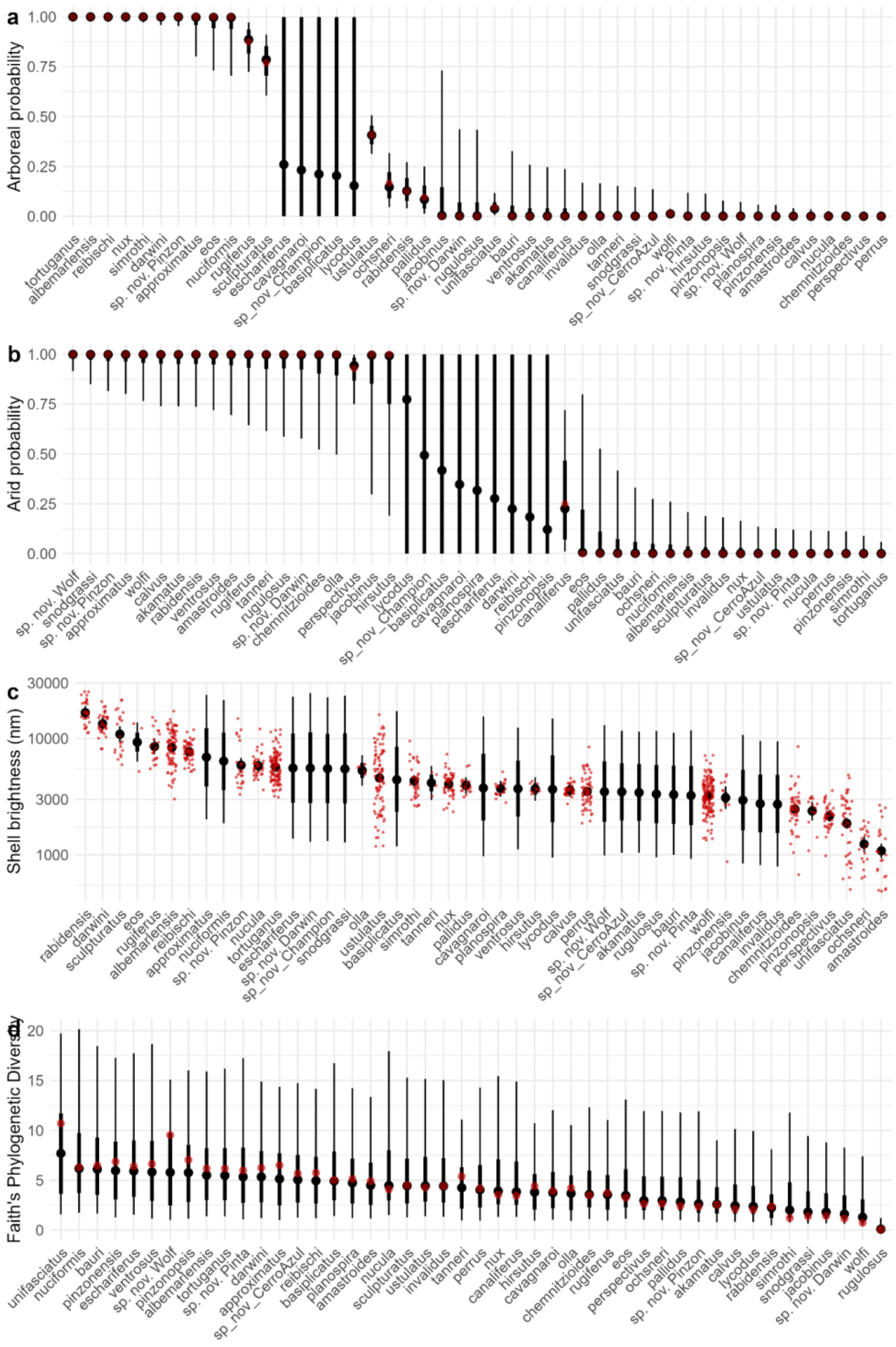


**Figure S8. Model predictions for nematode diversity (Faith's PD) (Model Equation S1.2).**

Red points show the raw data; black points show model predictions (posterior means with 95% credible intervals).

### S1.3 Relationship between host speciation rate and network position

In host-parasite systems, host diversification and ecological shifts can reshape parasite associations through processes like host switching and range expansions (Hoberg & Brooks 2008). These models propose that periods of ecological opportunity allow host lineages to expand into new niches, potentially escaping ancestral parasites via shifts in habitat, behavior, or physiology. In host-parasite systems, such expansions can lead to temporary parasite reduction or novel associations. Applying these ideas to the *Naesiotus* radiation, we hypothesize that rapidly diversifying lineages, particularly those adapting to arid environments or arboreal niches, may have evaded interaction with certain nematode clades. If so, diversification would leave a detectable footprint on network structure, with recently speciated hosts occupying more peripheral positions.

To test this, we estimated speciation rates across the *Naesiotus* radiation using ClaDS (Cladogenetic Diversification Rate Shift) (Maliet et a. 2019), and computed their network centrality as the first axis of a PCA composed of three complementary metrics: degree, betweenness, and eigen centrality. The final dataset included 40 *Naesiotus* species, as 7 species from the 47 present in the network were excluded from the analysis. Six of these were not included in the phylogeny: *pinzonopsis*, which has not yet been assessed, and five species considered extinct (*cavagnaroi*, *eschariferus*, *lycodus*, *olla*, and *planospira*). One additional species, *sp. nov. Champion*, was excluded because we were unable to amplify nematode DNA from its encapsulations.

We then fitted phylogenetic mixed models to test whether species' centrality in the host-parasite network was associated with their speciation rates. The response variable was the principal component summarizing centrality (from the PCA), and predictors included the mean speciation rate (estimated from ClaDS), island age (minimum emergence), log-transformed island area, and guild size (G), defined as the number of *Naesiotus* species co-occurring on the same island. All predictors were standardized (mean = 0, SD = 1) prior to analysis.

To propagate uncertainty in speciation rate estimates, we generated 100 replicate models. In each replicate, one value was sampled from the posterior distribution of ClaDS speciation rates for each species, and the model was fit to these sampled rates. Species identity was included as a random effect with a phylogenetic covariance structure derived from the *Naesiotus* tree to account for shared evolutionary history:

$$PCAcentrality \sim \beta_0 + \beta\lambda_s + \beta_{age}Age_i + \beta_{area}\log(Area_i) + \beta_G G_i + \alpha_s^{phylo} + \epsilon_i, \qquad \text{Equation S1.7}$$

$$\alpha^{phylo} \sim MVN(0, \sigma^2_{phylo}A)$$

Where $\lambda_s$ is the speciation rate for a species *s* sampled from the posterior predicted by ClaDS, and *A* is the phylogenetic relatedness matrix. Each model was implemented using the *MCMCglmm* package with a Gaussian error distribution and weakly informative priors on both the residual and random components. Models ran for 1,000,000 iterations with a burn-in of 500,000 and thinning interval of 100.

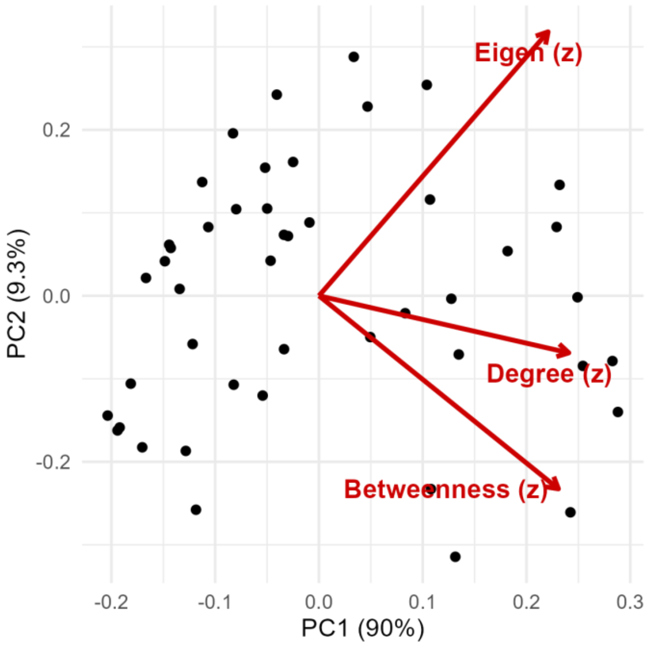


**Figure S9. Principal component analysis (PCA) of three centrality measures for *Naesiotus* species within the regional network.**

The first principal component (PC1) was used as the response variable in the model.

### S1.4 Regional interaction network

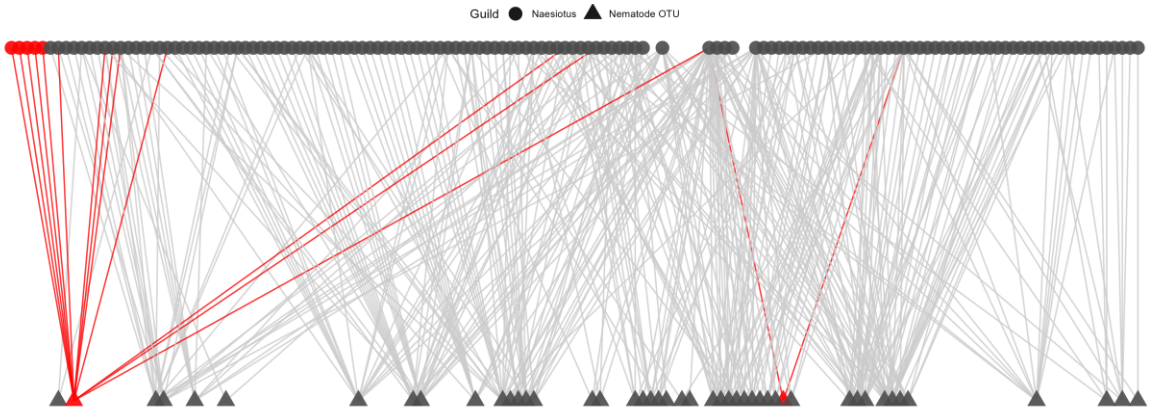


**Figure S10. Archipelago Naesiotus-nematode network.**

The network show 46 *Naesiotus* species (triangles) and 147 nematode OTUs (circles). Naesiotus species thought to be extinct are shown in red. Nematode OTUs shown in red interact only with these presumably extinct *Naesiotus*, indicating potential secondary extinctions.

### S1.5 Phylogenetic signal of nematode load and diversity

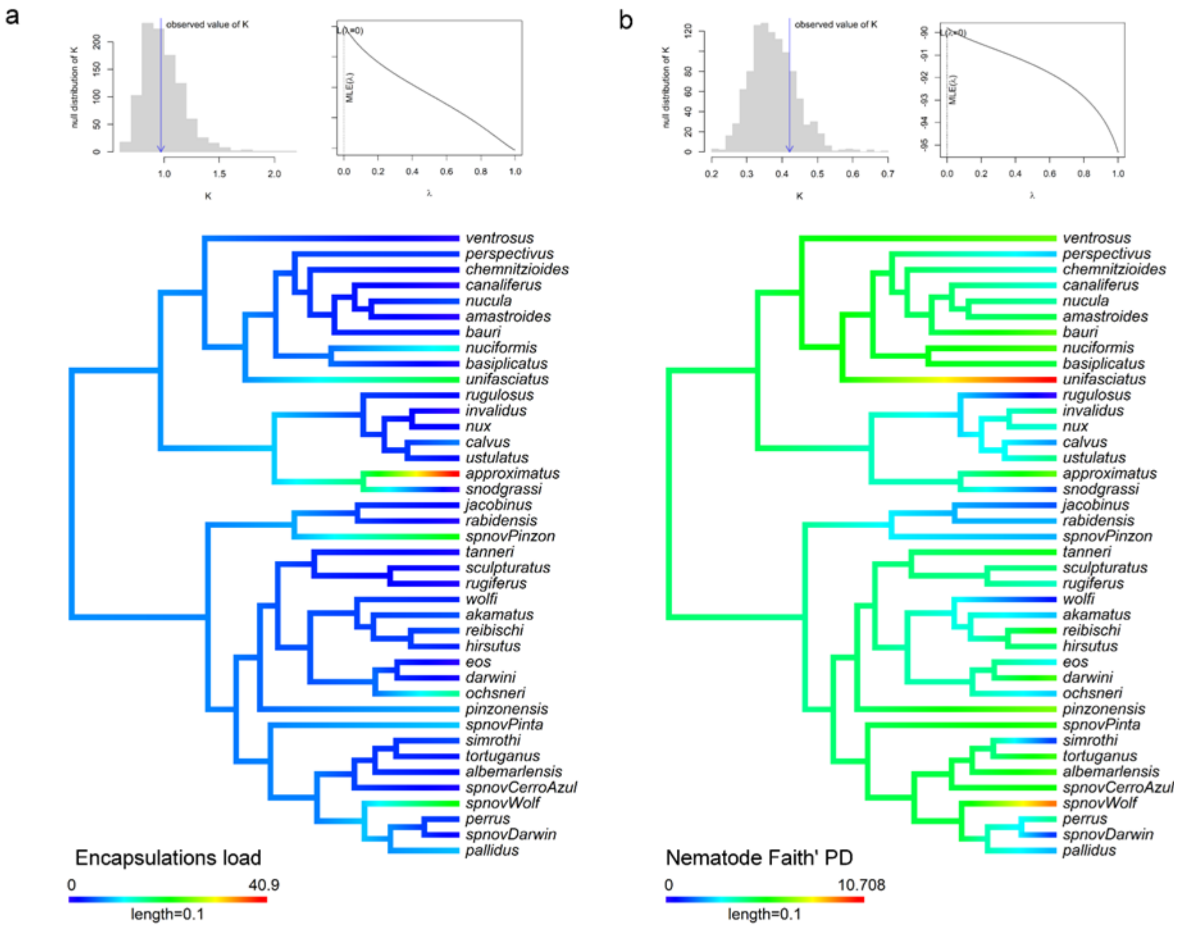


**Figure S11. Phylogenetic signal in nematode encapsulation load and diversity.**

Cropped trees of the *Naesiotus* radiation showing nematode encapsulation (a) load and (b) diversity, visualized as continuous trait maps. The associated graphs display the results of Blomberg's K and Pagel's lambda tests for phylogenetic signal.
